# Transplantation of Endothelial Progenitor Cells Solely Leads to Development of Functional Neo-vessels *in vivo*

**DOI:** 10.1101/049650

**Authors:** Rokhsareh Rohban, Nathalie Etchart, Thomas R. Pieber

**Affiliations:** Department of Internal Medicine, Division of Endocrinology and Diabetology, Medical University of Graz, Austria; Center for Medical Research (ZMF), Medical University of Graz, Austria

**Author notes:** Correspondence: Rokhsareh Rohban, PhD and Thomas R. Pieber, MD Division of Endocrinology and Diabetology, Medical University of Graz, Austria.

## Abstract

It has been believed that de novo vessel formation (neo-vasculogenesis) can be induced by co-transplantation of pericytes or mesenchymal stem/progenitor cells (MSPC) with endothelial cells or endothelial colony-forming cells (ECFC). The requirement for co-transplantation of two adult progenitor cells is one factor that can potentially complicate the process of therapeutic vasculogenesis which hampers the development of strategies for therapeutic intervention referred to as ‘regenerative medicine’. Here we employed a novel strategy for therapeutic vessel development by transplanting endothelial colony forming progenitor cells solely to immune compromised mice and detect vessel formation capacity of single ECFC transplants compared to ECFC/MSPC co-transplants. We applied umbilical cord derived and bone marrow derived-MSPC and umbilical cord derived ECFC with different total cell number for subcutaneous transplantation in matrix composites either alone or mixed at a ratio of 1:5 subcutaneously into immune deficient NOD.Cg-Prkdc^scid^ Il2rg^tm1Wjl^/SzJ; NSG mice. Implants were harvested one day, one, two, eight and 24 weeks after transplantation for detecting the state of vessel formation and stability of the transplants by histological assessments. Additionally, endothelial progenitor cells derived from various human tissues such as umbilical cord blood, peripheral blood and white adipose tissue were used to assess their potential for vessel formation *in vivo*.

Results confirmed that single transplantation of ECFCs with a higher cell number and later in the time course after transplantation is as efficient as co-transplantation of ECFC with MSPC at forming stable-perfused human vessels. Amongst ECFCs isolated from different human sources, white adipose tissue derived ECFC are most potent in forming neo-vessels (micro-vessels) in vivo, thus WAT-ECFC could be an optimal cell for vasculogenesis regenerative application.

Co-transplantation of ECFC and MSPC with the defined 5:1 ratio or sole ECFC with a higher cell dosage was essential for vessel generation *in vivo*.

## 1. Introduction

neo-vessel formation is a crucial step that takes place in tissue regeneration as well as malignancies like tumor formation which is mainly associated with angiogenesis in adult life (Carmeliet and Jain, 2011).The formation of stable, functional vasculatures by stem and progenitor cells is referred to as neo-vasculogenesis. The procedure initiates through migration of endothelial (progenitor) cells forming the inner layer of newly formed neo-vessels and mesenchymal cells serving as pericytes maintaining vasculature stability (Reinisch et al., 2007, Losordo and Dimmeler, 2004, Carmeliet and Jain, 2011). There are evidences through studies on neo-vessel formation that suggest establishing of neo-vessels does not exclusively require involvement of pericytes as micro-vessel supporter e.g. in the course of vessel formation and vasculature expansion from endothelial cells in tumors (REF). In development phase of embryo, vessel formation occurs in alignment with re-arrangement circumstances through which the development of new vasculatures and pruning of others takes place (Risau and Flamme, 1995). Vessel formation takes place presumably upon the interaction of cellular secreted factors including various signaling components (Dasari et al., 2010, Dumont et al., 1992, Rohban et al., 2013). We used an established vessel formation model (Au et al., 2008, Mead et al., 2008, Reinisch et al., 2009) to test our hypothesis of whether neo-vasculogenesis requires the combination of ECFC and MSPC or can be performed only through ECFC transplantation. We also aimed to investigate whether the neo-vessels derived from ECFC are stable and functional in a longer time period. We further tested the vasculogenesis potential of ECFCs isolated from various human tissues such as UCB, UC, PB and WAT. Here we show that increased number of transplanted ECFC can result in perfused neo-vessel formation in an extended time course *in vivo*. The ECFC derived micro-vessels are functional and stable *in vivo* for up to 6 months, and WAT-ECFC is the most potent ECFC source in terms of neo-vasculogenesis. This data suggests a new regenerative vasculogenesis strategy that can be used during experimental neo-vessel formation *in vivo*.

## 2. Materials and Methods

### 2.1. Ethics Statement

Human tissue samples were collected upon written informed consent from healthy individuals according to procedures approved by the Ethical Committee of the Medical University of Graz (Protocols 19-252 ex 07/08, 18-243 ex 06/07, 21.060 ex 09/10, 19-252 ex 07/08). Adult tissue samples were obtained after receiving written informed consent from healthy donors. Umbilical cord (UC) and umbilical cord blood (UCB) samples were collected following full-term deliveries with informed consent from the mothers according to the Declaration of Helsinki.

All animal experiments were carried out according to the 2010/63/EU guidance of the European Parliament on the welfare of laboratory animals. Protocols obtained the approval of the Animal Care and Use Committee of the Veterinary University of Vienna on behalf of the Austrian Ministry of Science and Research.

### 2.2. Preparation of ECFC and MSPC Culture Medium

ECFC isolation and expansion was carried out using endothelial basal medium (EBM-2) supplemented with 100 μg/mL Streptomycin, 100 U/mL penicillin, 2 mM L-Glutamine (all Sigma) as well as manufacturer-provided aliquots containing vascular endothelial cell growth factor (VEGF), human epidermal growth factor (EGF), insulin-like growth factor-1 (IGF-1) and human basic fibroblast growth factor (bFGF). The medium was also supplemented with 10% pooled human platelet lysate (pHPL) replacing fetal bovine serum (FBS) as described previously (Reinisch et al., 2007, Schallmoser et al., 2007b, Reinisch et al., 2009, Hofmann et al., 2012b). Briefly, an amount of 450 ± 45 mL of the blood was collected according to the informed consent from healthy individuals following the Austrian protocol for blood donation. In order to produce platelet enriched plasma, the buffy coat was subjected to red blood cell and plasma separation procedures automatically and thereafter transferred into satellite containers (Compomat, G3, NPBI, Amsterdam, Netherlands). Platelet-rich plasma (PRP) was further exposed to leukocyte depletion infiltration procedures (PALL Autostop, Pall, Dreieich, Germany) and thereafter stored at −30°C to allow for platelet cleavage. To avoid the incidence of individual donor variations, a minimum number of 10 platelet lysate units were pooled and stored at −30°C until further use (Schallmoser et al., 2007a, Schallmoser and Strunk, 2009).

Prior to pHPL supplementation in the complete media, an amount of 10 U/mL of Heparin (Biochrom AG) was added to avoid pHPL solidification within the liquid medium.

MSPC isolation and expansion were performed using α-modified minimum essential medium (α-MEM, M4526; Sigma-Aldrich; St. Louis; MO, USA; www.sigmaaldrich.com) supplemented with 100 μg/mL Streptomycin as well as 100 U/mL Penicillin (to minimize the bacterial contamination), 2 mM L-Glutamine (all Sigma;) and 10% pHPL (Schallmoser et al., 2007a, Bartmann et al., 2007, Schallmoser and Strunk, 2009). To avoid platelet lysate solidification in the medium, heparin (2 U/mL, Biochrom AG) was added to the α-MEM medium prior to pHPL supplementation.

### 2.3. ECFC and MSPC Isolation and Expansion

MSPC were isolated and expanded from umbilical cord and bone marrow (Shahdadfar et al., 2005, Szoke et al., 2012, Kita et al., 2010, Miao et al., 2006).

Umbilical cord-derived MSPC were isolated from Wharton’s jelly dissected from umbilical cord and cut into pieces (4-6 mm^2^). The tissue pieces were placed on the culture plate to allow for plastic attachment prior to addition of the culture medium (http://www.jove.com/author/AndreasReinisch). The cells were allowed to grow for 2-3 weeks.

ECFC were isolated and expanded from human umbilical cord, umbilical cord blood, adult peripheral blood and white adipose tissue (Prasain et al., 2012, Hofmann et al., 2009, Reinisch and Strunk, 2009, Mead et al., 2008, Szoke et al., 2012). For ECFC isolation from blood, cell culture was initiated no more than two hours after blood donation. To minimize cell loss, any unnecessary manipulation of freshly collected blood including red blood cell lysis or density gradient centrifugation was avoided. The heparin-containing blood was diluted in EGM-2 in a ratio of 1:4. The cultures were incubated at 37°C, 5% CO_2_ overnight and washed twice with pre-warmed PBS to remove non-adherent cells prior to addition of the fresh pre-warmed EGM-2 medium. The medium was replaced 2-3 times per week until the cobblestone-shaped ECFC colonies appeared (Reinisch et al., 2007, Schallmoser et al., 2007b, Reinisch et al., 2009, Hofmann et al., 2012b).

In order to expand ECFC and MSPC, the frozen cells of the first passage were thawed and seeded in 2,528 cm^2^ cell factories (CF-4, Thermo Fisher Scientific, Freemont, CA) using 500 mL EGM-2 and α-MEM medium, respectively. The cell-containing CF4s were maintained at 37°C, 5% CO_2_ and 95% humidity in an incubator. The cultures were subjected to medium replacement twice weekly to allow for the required nutrition accessibility and removal of the waste. Upon reaching 70-80% confluence, the cells were harvested using pre-warmed trypsin (50 mL per CF-4, 4-5 min., 37°C; Sigma Aldrich). After two washes with pre-warmed PBS and centrifugation (5 min, 300xg, 4°C), the harvested cells were quantified and checked for viability by means of Bürker Türk hemocytometer and trypan blue exclusion, respectively.

### 2.4. Flow Cytometric Characterization of ECFC and MSPC

ECFC and MSPC were stained with cell surface markers and the reactivity was evaluated using flow cytometry (Facs Calibur Flow cytometer, BD) according to the manufacturer’s instructions (see Fig‥‥). The ECFC and MSPC cellular populations were checked for purity using marker profiling prior to use for *in vitro* and *in vivo* experiments Accordingly, trypsinized cell populations were considered pure when they were ≥95% positive for their positive markers and ≤2% positive for their negative markers (Dominici et al., 2006). MSPC and ECFC populations were thereafter stored long term to be used for *in vitro* and *in vivo* assays.

### 2.5. Animal Experiments

Animal experiments in this project were performed following the 2010/63/EU guidance of the European Parliament on the welfare of laboratory animals. Protocols were approved by the Animal Care and Use Committee of the Veterinary University of Vienna on behalf of the Austrian Ministry of Science and Research. Immune-deficient NOD.Cg-Prkdc^scid^ Il2rg^tm1Wjl^/SzJ (NSG) mice were purchased from the Jackson laboratory (Bar Harbor, ME, USA), housed in specific pathogen-free (SPF) facility of the Medical University of Graz and were used for the *in vivo* experiments between seven to 18 weeks of age. The immune-compromised NSG mice were used in order to avoid the immune response leading to the rejection of injected human cell transplants. ECFC/MSPC or ECFC alone were re-suspended with a ratio of 80:20 % in ice-cold extracellular matrix (300 mL per plug, Cat. No. ECM 625, Millipore, Billerica, MA, USA), thereafter injected in form of subcutaneous plugs into the flank of the NSG mice (Melero-Martin et al., 2007, Au et al., 2008). Prior to injection, mice were anesthetized following the approval for animal handling (BMWF:-66.010/0082-II/10b/2009).

Mice were observed for the indicated time periods of two to 24 weeks after injection. At the defined time point, the mice were sacrificed by cervical dislocation and the plugs were surgically dissected from the subcutaneous tissue, photographed using stereomicroscope (SZX12, Olympus) and fixed in neutral buffered 4% paraformaldehyde overnight to be used for histological assessments.

### 2.6. *In vitro* capillary-like structure formation

Umbilical cord blood, umbilical cord, white adipose tissue and peripheral blood-derived ECFC were seeded in 225 cm^2^ tissue culture flasks (BD Biosciences) and incubated in 37°C until confluence. Upon trypsinization followed by a washing step with pre-warmed PBS, 7.5 x 10^4^ ECFC were re-suspended in 2 mL EGM-2 supplemented with 10% pHPL and seeded on 9.2 cm^2^ polymerized Matrix (Angiogenesis assay kit; Millipore, Billerica, MA, USA) as described previously (Rohde et al., 2007). After 24 hours, endothelial networks were captured using a Color View III camera on an Olympus IX51 microscope with the analySIS B acquisition software (all Olympus, Hamburg, Germany). Numbers of branching points were evaluated using ImageJ software (http://rsbweb.nih.gov) as described (Hofmann et al., 2012a).

### 2.7. Vessel Formation *In Vivo*

ECFC and MSPC were subjected to an optimized isolation and purification procedure as previously indicated (Reinisch and Strunk, 2009).ECFC were seeded in EGM-2 (Lonza) at a density of 1,000 cells/cm^2^ and MSPC in α-MEM (Sigma-Aldrich, St. Louis, MO) at a density of 500 cells/cm^2^ in 2,528 cm^2^ cell factories (CF-4, Thermo Fisher Scientific, Freemont, CA).

In a different set of *in vivo* vascularization experiments, a total number of 2x10^6^, 8x10^6^ or 32x10^6^ cells were used to investigate vessel formation. Two, eight or 32 million MSPC (MSPC only), two, eight or 32 million ECFC (ECFC only) or the combination of 1.6 x 10^6^, 6.4x10^6^ or 24x10^6^ ECFC with 0.4 x 10^6^, 1.6x10^6^ or 8x10^6^ MSPC (ECFC+MSPC) were re-suspended in 300 μL ice-cold extracellular matrix (*In vitro* angiogenesis assay kit, Millipore, Billerica, MA, USA) and injected subcutaneously into immune-deficient NSG mice (four plugs per mouse were injected, for every cell condition three mice were used). For every set of *in vivo* experiments, cell-free ‘Matrix only’ plugs (Millipore) were injected and used as controls.

To investigate the variability of neo-vasculogenesis potential amongst ECFCs from different sources, a total number of 2x10^6^ ECFCs from UC, UCB, PB and WAT were resuspended with 300 mL Matrix (In vitro angiogenesis assay kit, Millipore) and injected to NSG mouse subcutaneously. The plugs were surgically removed after 2 weeks and the plugs were stored in Formaldehyde (‥‥…) to be used for histological experiments.

Mice were sacrificed at day one, 14, 56 or 168 after implantation, and plugs were harvested from the subcutaneous sites (three mice and three plugs per cell combination per time point).Plugs explanted after one day were subjected to IHC and tunnel assay to investigate the number (rate) of apoptotic cells,whereas parallel plugs explanted at two, eight and 24 weeks (14, 56 and 168 days, respectively) were used for histological experiments for confirmation of functional and stable vessel formation in a time-course analysis.

### 2.8. Histological Experiments and Analysis

Plugs harvested after 2, 8, and 24 weeks were fixed using 3.7% neutral-buffered formalin in 4°C overnight thereafter dehydrated in a graded series of ethanol prior to being embedded in paraffin. Paraffin-embedded tissue sections (4μm) were deparaffinized using xylene and descending alcohol series prior to Hematoxylin/Eosin (H&E), Movat’s pentachrome as well as immune histochemistry staining.

#### Hematoxylin-Eosin Staining (H&E)

Sections (4 μm) of formalin-fixed (3.7% neutral buffered, overnight) paraffin-embedded plugs were deparaffinized using heat (68°C hot chamber) for an hour followed by treatment with xylene and descending alcohol series (5 min each) and stained by routine hematoxylin-eosin (H&E) protocol (Lillie, 1965).The sections were stained using Mayer’s hematoxylin (Sigma Aldrich) for 3 min, thereafter rinsed in running tap water and consequently stained with eosin phloxin (Sigma Aldrich) for 30 sec. Sections were then washed with running water, rehydrated in ascending alcohol series (2 min each) and mounted using TissueTek^®^ Glas ™ mounting media (Sakura). The nuclei, cytoplasm and red blood cells were differentially stained blue, purple and red, respectively (Lillie, 1965). The protocol has been optimized based on the specimen properties.

To quantify vessel formation, luminal structures containing red blood cells were considered as perfused micro-vessels and counted using ImageJ (http://rsbweb.nih.gov). For each condition, 5 high power fields (200x original magnification) of H&E-stained slides were examined by two independent observers using counting function in ImageJ software. For *in vivo* vasculogenesis experiments, micro-vessel density was quantified by analyzing the number of red blood cell-containing luminal structures additionally at three different cutting depths (150 μm intervals).

#### Immune Histochemistry Staining

Sections (1.5 μm) of formalin-fixed (3.7% neutral buffered, overnight) paraffin-embedded plugs were de-waxed prior to antigen retrieval using heat (68°C/160 W, 40 min) at either high or low pH (pH 9 or 6, respectively) depending on the protein target and antibody properties followed by a descending alcohol series (2x xylol, 5 min each; 1x ethanol 100%, 5 min; 1x ethanol 90%, 5 min; 1x ethanol 70%, 5 min; 1x ethanol 50%, 5 min; 1x PBS, 5 min, distilled water, 5 min) as previously described (Hofmann et al., 2012a). Endogenous peroxidases were inhibited using H_2_O_2_ (15 min) and non-specific antibody binding was minimized using Ultra V Block (Thermo Scientific; 7-10 min), mouse-on-mouse blocking (MOM, Vector; 90 min) and serum-free protein inhibiting reagent (Dako; 30 min). Sections were incubated (45 min, RT) with unconjugated monoclonal mouse anti-human antibodies against CD31 (clone: JC70A, 5.15 μg/mL, Dako), CD90 (clone: EPR3132, Abcam, Cambridge, MA, USA) Vimentin (clone: V9, 0.78 μg/mL, Dako), von Willbrand factor (vWF, clone F8/86, Dako) or appropriate amount of IgG1 (BD) as the control. The signal was developed using Ultravision LP large volume detection system horseradish peroxidase (HRP) polymer (Thermo Scientific) followed by diaminobenzidine (DAB) or alkaline phosphatase detection system using either fast blue or fast red (Vector) according to the manufacturer’s protocols. Avidin-biotin inhibiting (Vector) was used before staining with biotinylated polyclonal rabbit anti-human CD90 (EPR3132, Abcam) and biotinylated monoclonal goat anti-rabbit IgG1 (BD). Streptavidin-horse radish peroxidase conjugate (Dako) and diaminobenzidine (DAB) were used to detect and visualize the positive signal. Sections were counter-stained using hematoxylin for 1-2 min. The protocol has been optimized based on the specimen and antibodies properties.

In order to quantify vasculogenesis in ECFC+MSPC plugs and ECFC only explants or plugs, luminal structures filled with RBC representing hu-CD31 and vimentin positive signal were considered as perfused functional vessels and quantified by two independent observers using five high power microscopic fields (200x original magnification) of H&E-stained sections from the related plugs using ImageJ software. The number of perfused vessels was quantified using ImageJ software (http://rsbweb.nih.gov).

#### Movat’s Pentachrome Staining

Paraffinized sections (4 μm) were hydrated using distilled water and stained in alcian blue (stain for ground substance and mucin, Dako, 20 min). The slides were washed under running water for 10 min followed by alkaline alcohol treatment (to convert the alcian blue into insoluble monastral fast blue) for 1-2h and washing steps (10 min) to remove alkaline alcohol. The sections were incubated with Verheoff’s hematoxylin solution (15 min, Sigma Aldrich) to promote nuclei and elastic fiber staining. Upon rinsing with dH_2_O (4 times, each time for 2 min), the slides were exposed to 2% aqueous ferric chloride several times to promote differential staining of the different tissues. The differentiation process was checked microscopically for black elastic fiber staining versus gray background. Upon washing steps using sodium thiosulfate (1 min) and distilled water (5 min), the sections were stained with scarlet-acid fuchsine (8:2, Dako) for 90 sec (Stain for fibrinoid, fibrin-intense red, muscle-red). After rinsing steps in distilled water and 0.5% acetic acid water, the slides were incubated with 5% aqueous phosphotungstic acid (5-10 min) and checked microscopically for further differentiation to collagen indicator (pale pink) versus bluish ground substance. After differentiation was completed, the slides were rinsed in 0.5% acetic acid water (5 min, 2X) followed by 100% alcohol (5 min, 3X). (Color code here!)

### 2.9. Tunnel Assay

In order to test apoptosis *in vivo* and *in vitro*, ECFC+MSPC plugs and ECFC only plugs harvested one, 14 and 56 days post transplantation and ECFC+MSPC admixed with matrix incubated at 37°C for three and five days *in vitro* were analyzed for apoptosis using TdT-mediated dUTP nick end labeling (TUNEL Assay; Dead End^TM^ Fluorometric System, Promega) in accordance with manufacturer’s instructions. Nuclei were counter-stained using propidium iodide. To develop a positive control, slides were incubated with 1U DNase 1 (10 min, RT, Fermentas). For a negative control, incubation buffer was prepared without rTdT. Signals were documented using confocal microscopy (LSM510 Meta, Zeiss).

### 2.10. Statistics

All data are shown as mean ± standard deviation (SD). Statistical differences were determined using unpaired student’s t-test or ANOVA using post-hoc Bonferroni test for multiple comparisons. Differences were considered significant (*) when the P-value was less than 0.05, very significant (**) when P-value was less than 0.001 and extremely significant (***) when P-value was less than 0.0001.

## 3. Results

### 3.1. “ECFC only” population is capable of neo-vasculogenesis, independent of the MSPC population admix.

Subcutaneous transplantation of ECFC+MSPC in an established 80: 20 ratio (Reinisch et al., 2009) or ECFC population solely was carried out to perform a comparative study on neo-vasculogenesis potential of these mesodermal progenitor cell types (combinations) *in vivo*.

Plugs containing UC-MSPC+UC-ECFC and BM-MSPC+UC-ECFC were transplanted as two pre-vasculogenesis transplantation models as previously described (Rohban et al., 2013). The use of two different source of isolation for the MSPC admix was to compare the impact of UC-and BM-MSPC in contributing to neo-vessel formation.

UC-ECFC solely was also transplanted to investigate the potential of vessel formation of ECFC single population. Upon plug harvesting after one, two, eight and 24 weeks (6 months), vessel formation was evaluated macroscopically and histologically in the ECFC+MSPC and “ECFC only” explants containing 2, 8 and 32x10^6^ cells, respectively (three mice and three implants per condition and time point were used).Macroscopic features of the explanted plugs indicated the development of perfused red blood cell-containing vessels in the plugs in vivo that was further confirmed by immune histochemistry analysis. The red color of the plugs explanted after one, two, eight and 24 weeks macroscopically indicated patent red blood cell-containing mature micro-vessel formation after ECFC+MSPC, as well as sole ECFC transplantation (Figure).

The result confirmed the ability of MSPC/ECFC to develop micro-vessels regardless of the source of MSPC admixed. However, a slight/significant difference in the number of created micro-vessels is detectable (Figure). The result also showed that sole ECFC transplants were able to vascularize *in vivo* (Figure), the number of created vessels within the ECFC single transplants was not significantly hampered compared to ECFC/MSPC co-transplants (Figure).

Human vimentin immunehistochemistry staining indicates that the created vessels are originated from human mesodermal derived stem and progenitor cells (Fig……).

### 3.2. Vessels created by ECFC maintain long lasting stability and functionality in vivo.

Stability and functionality of the created micro-vessels where investigated in a later time point using penta chrome staining. Formation of elastic fibers in the outer layer of neo-vessels is distinguished by brown-black color in Penta-chrome staining method. As seen in Fig…, both vessels created in co-transplants and ECFC single transplants have developed a layer of elastic fibers in the outer layer, indicating the stability of the created vessels in both transplants late in the time point (24 weeks). The newly formed vessels are filled with RBC presumably as a result of connecting to the mouse circulation, which indicates the functionality of the neo-vessels in vivo. The data revealed that the vessels created by “ECFC only” are long lasting and functional in vivo.

### 3.3. ECFC associated vasculogenesis differentiation is dependent on the injected cell dose and time of incubation in vivo.

MSPC/ECFC and ECFC were transplanted with three different total cell numbers at a ratio of 20:80 and 0:100, respectively. Co-transplants containing 2x10^6^, 8x10^6^ and 32x10^6^ cells were compared to that of ECFC single transplants with regard to the number of created vessels at different time points in vivo. The result showed that vessel formation in co-transplants containing 2x10^6^ cells is detectable after one week, whereas ECFC single transplants containing 2x10^6^ cells are vascularized eight weeks post transplantation (Figure). Both transplants remained vascularized for up to six months after transplantation (Figure). In transplants with a fourfold increase in the number of implanted cells (8x10^6^), vasculogenesis in ECFC single transplants takes place two weeks after implantation (Figure), whereas in transplants with a total number of 32x10^6^ cells, neo-vessels are formed in ECFC only transplants one week post implantation (Figure). This data suggests that the vessel formation in ECFC only transplants depends on the number of transplanted cells (Fig) as well as in vivo incubation time course (Figure).

### 3.4. Amongst ECFC of various tissue sources, WAT-ECFC were shown to have the highest potential of neo-vasculogenesis in vivo.

ECFC were isolated from various human tissues such as umblical cord (UC), umbilical cord blood (UCB), peripheral blood (PB) and white adipose tissue (WAT), and the cells were transplanted at a density of 2x10^6^ and 8x10^6^ to the NSG mice, subcutaneously. Upon plug removal after two weeks, the number of vessels was quantified using histological techniques. The result revealed that amongst ECFCs derived from various tissues, WAT-ECFC are the only cells capable of vasculogenesis when transplanted at a density of 2x10^6^ in a two week time period, whereas nearly all ECFC sources were able to commit vasculogenesis upon transplantation at a density of 8x10^6^. This data unravels the high potential of WAT-ECFC to contribute in neo-vessel formation in vivo.

## 4. Discussion

De novo vessel formation (neo-vasculogenesis) is an essential step which takes place in course of organ regeneration, wound healing, inflammation as well as tumor growth (Segura et al., 2002, Elmore, 2007, Krysko and Vandenabeele, 2008). Vascular structure formation consists of migration and replication of endothelial colony forming cells (ECFC) as the backbone of newly formed vessel (Segura et al., 2002, Elmore, 2007, Krysko and Vandenabeele, 2008) and mesenchymal stem and progenitor cells (MSPC) as pericytes which serve as vessel supporters and maintain micro-vessel stability (Reinisch et al., 2007, Schallmoser et al., 2007b, Reinisch et al., 2009, Hofmann et al., 2012b). In the process of neo-vasculogenesis, the cross talk between the two cellular components through mediators and signaling molecules is inevitable, yet poorly understood. It is also not completely known whether single cell transplantation with manipulations in cell dose and implantation time course can lead in as successful vasculogenesis in vivo.

The aim of this study was to study the vasculogenesis potential of MSPC/ECFC co-transplantation compared to ECFC single transplantation. We also aimed to investigate whether various sources of ECFC differ in vasculogenesis commitment potential. The aim was accomplished by (1) histologically detecting the established micro-vessels in co-transplants and single transplant (2) quantifying and comparing the number of established neo-vessels in the transplants and (3) investigating the difference in vasculogenesis potential within various ECFC sources in vitro and in vivo.

We used a vasculogenesis model (Reinisch et al, 2009, and the others) to study the impact of different cell combinations in course of vessel formation in vivo. This model/strategy allowed us to compare the number of created vessels in transplants with different injected cell number and time course. Capillary like network formation (*In vitro* angiogenesis assay, Millipore) assessment and quantification of the number of branching points created by ECFC from different tissues helped to get an insight into different vasculogenesis potentials amongst ECFC isolated from various human tissues. In this study we compared vessel formation ability of the established MSPC/ECFC co-transplantation model with “ECFC only” transplant with different total number of cells and time points. It has been shown by Yoder et al. that the original cell that plays the major role in angiogenesis is endothelial cell and often circulating endothelial cells can enter a tomour and contribute to the profound angiogenesis process (ref). Medea et al. have also shown that endothelial progenitors can form hemangiomas in the brain tissue that are stable for a long period of time (REF). add the other papers supporting the idea e.g. Hofmann et al., 2013 (REFS). Contradictory earlier reports showed that vasculogenesis in vivo requires the co-transplantation of MSPC and ECFC as ECFC only transplantation cannot develop functional long lasting vessels in vivo, and the fragile layer of ECFCs need the support of MSPC or pericyte to stabilize the micro-vessel (REFS). Despite this ongoing controversy on the role/necessity of MSPCs in vasculogenesis processes, the precise role of MSPCs and the necessity of MSPC admix in course of neo-vasculogenesis has yet to be studied.

In our previous study, vessel formation was significantly hampered when ECFCs were treated with caspase-4 chemical blocker, suggesting the role of death associated signaling pathways and specifically inflammatory caspase-4 in vessel formation through single transplantation of ECFC (Rohban et al., 2013).

Our data also established the as of yet unknown role of ECFCs in covering the responsibility of MSPCs to stabilize the newly formed micro-vessels which has been shown to be dependent on the injected cell number and the duration of time course. In this study we compared the vasculogenesis potential of ECFCs isolated from various tissues, and distinguished the significantly higher potential of WET-ECFC for capillary like network formation in vitro as well as neo-vessel formation in vivo. WAT exists in adult body (REF) and can be dissected with rather low invasive methods (REF). ECFC isolated from WAT of individuals could serve as a patient-specific source of regeneration.

Unraveling the potential of ECFC single transplantation in establishing patent perfused neo-vessels with a high stability over time help us to develop better strategies for stem cell mediated tissue regeneration and will hopefully contribute to developing new therapeutic approaches. In addition we have unraveled a high vasculogenesis differentiation capacity of WET-ECFC, which can be attractive in terms of efficiency of cell sources to be used for cytotherapy and regeneration.

## 5. Conclusion

In this study the neo-vessel formation potential of MSPC/ECFC co-transplantation compared to ECFC single transplantation was assessed. “ECFC sole” associated vasculogenesis was shown to be highly dependent on the transplanted cell number as well as duration of in vivo incubation. Neo-vessels created by ECFC only population were shown to remain stable and functional for up to 6 months in vivo. ECFCs isolated from various tissues showed to possess variable capacity in terms of neo-vasculogenesis in vivo. WAT derived ECFCs showed to possess the highest potential in contributing to neo-vasculogenesis in vivo.

## Acknowledgements

The authors thank Birgit Feilhauer, Claudia Url and Daniela Thaler for their excellent technical assistance and Dr. Beate Boulgaropoulosfor structural proofreading and English language editing.

---

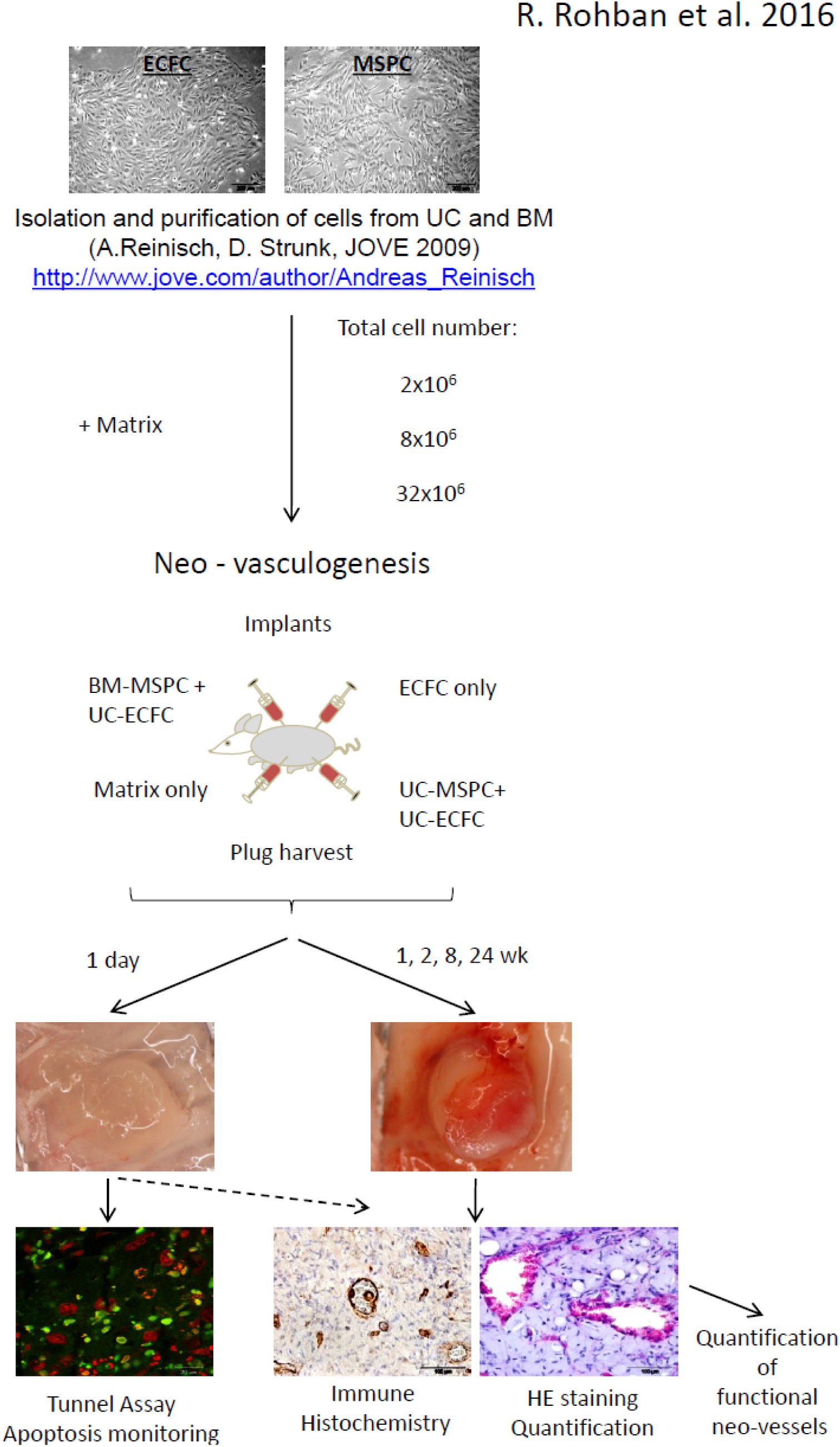

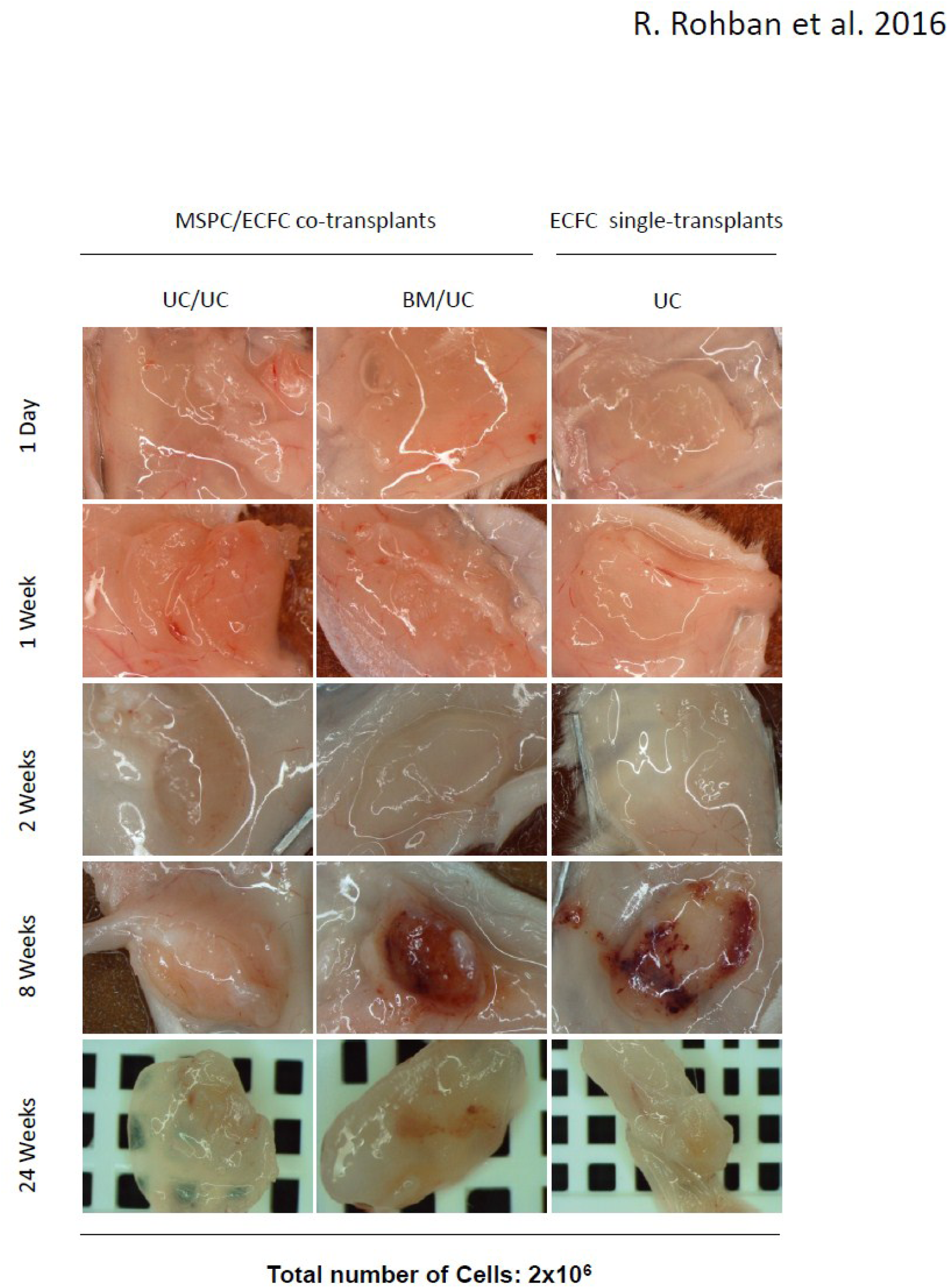

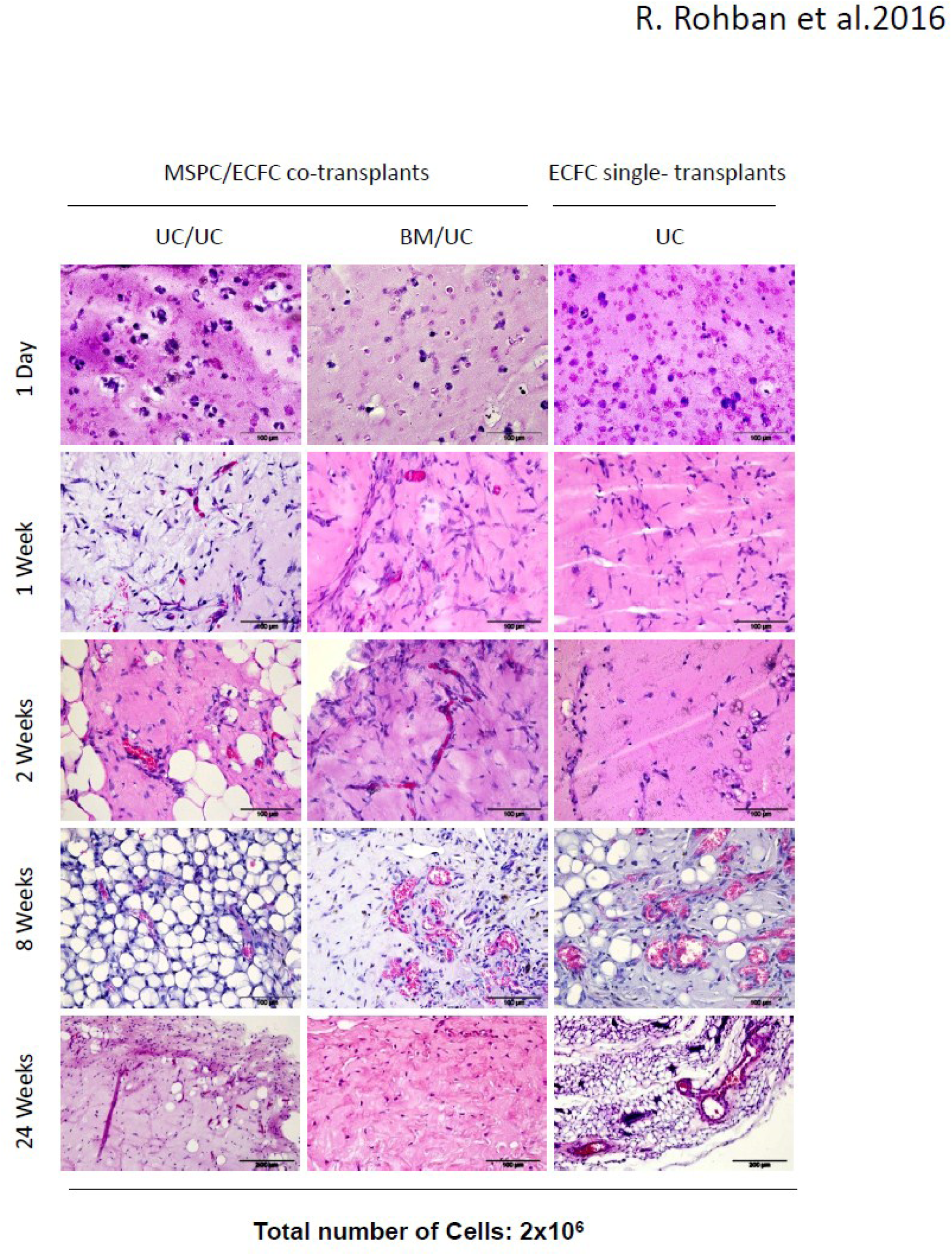

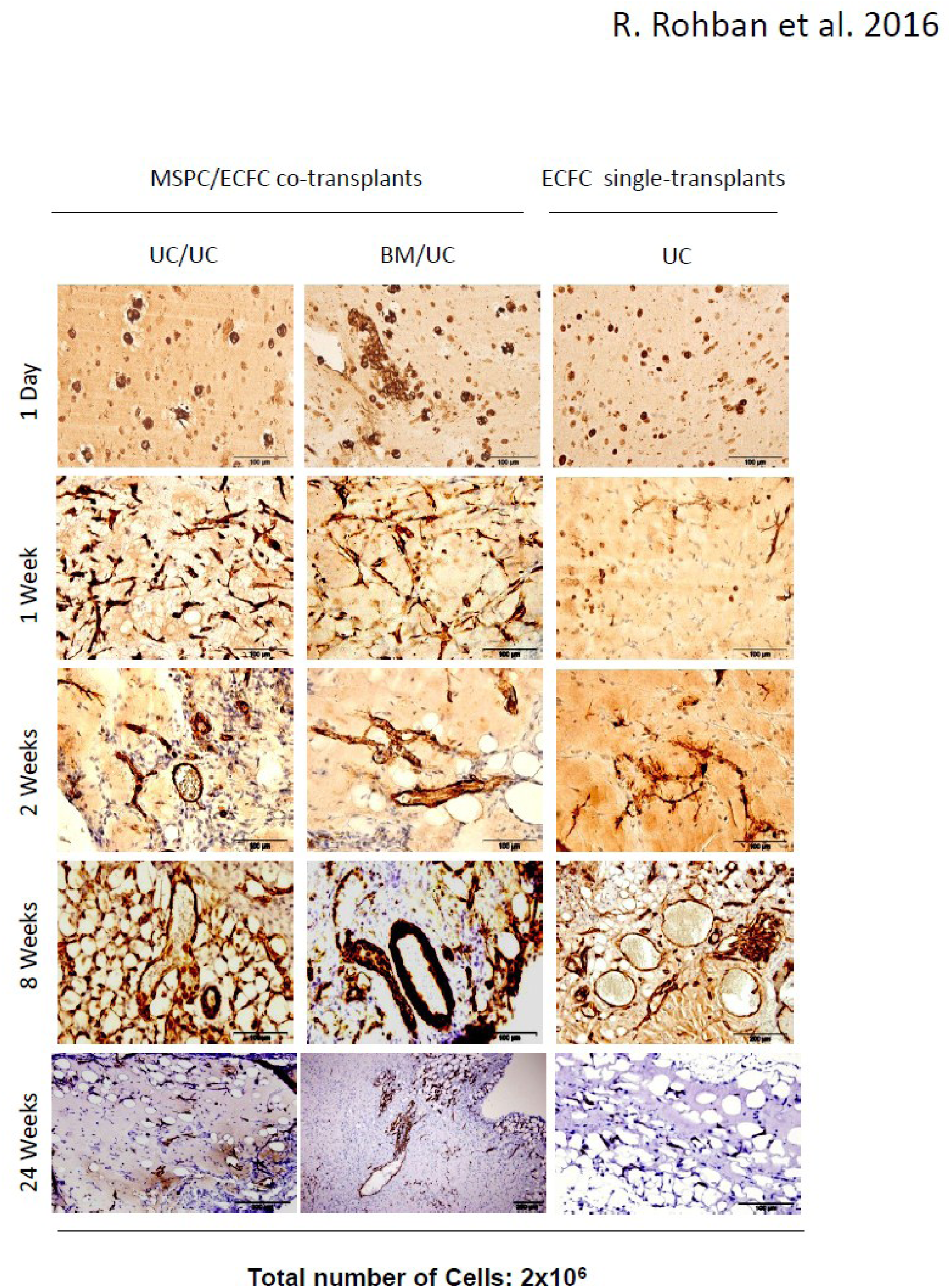

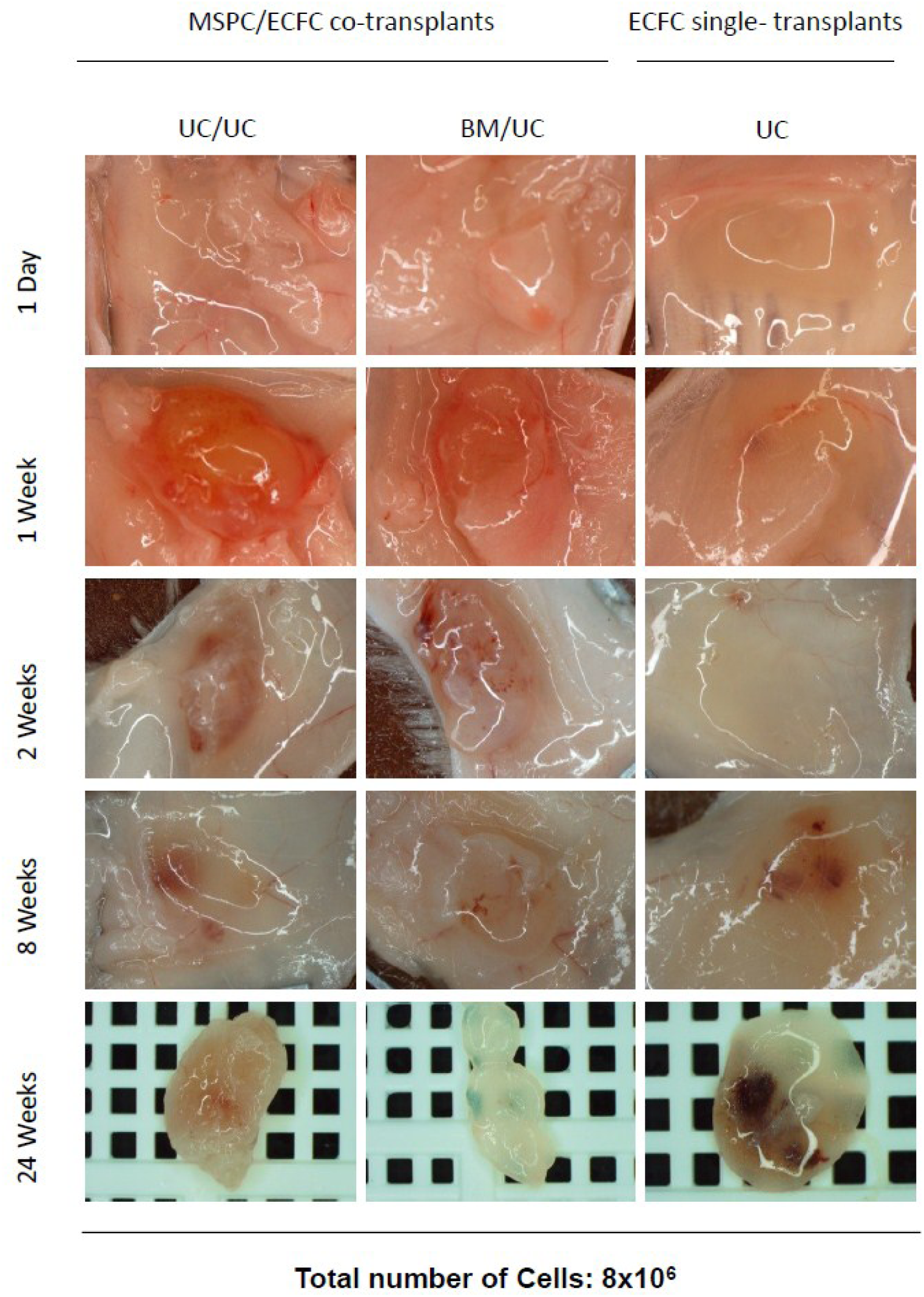

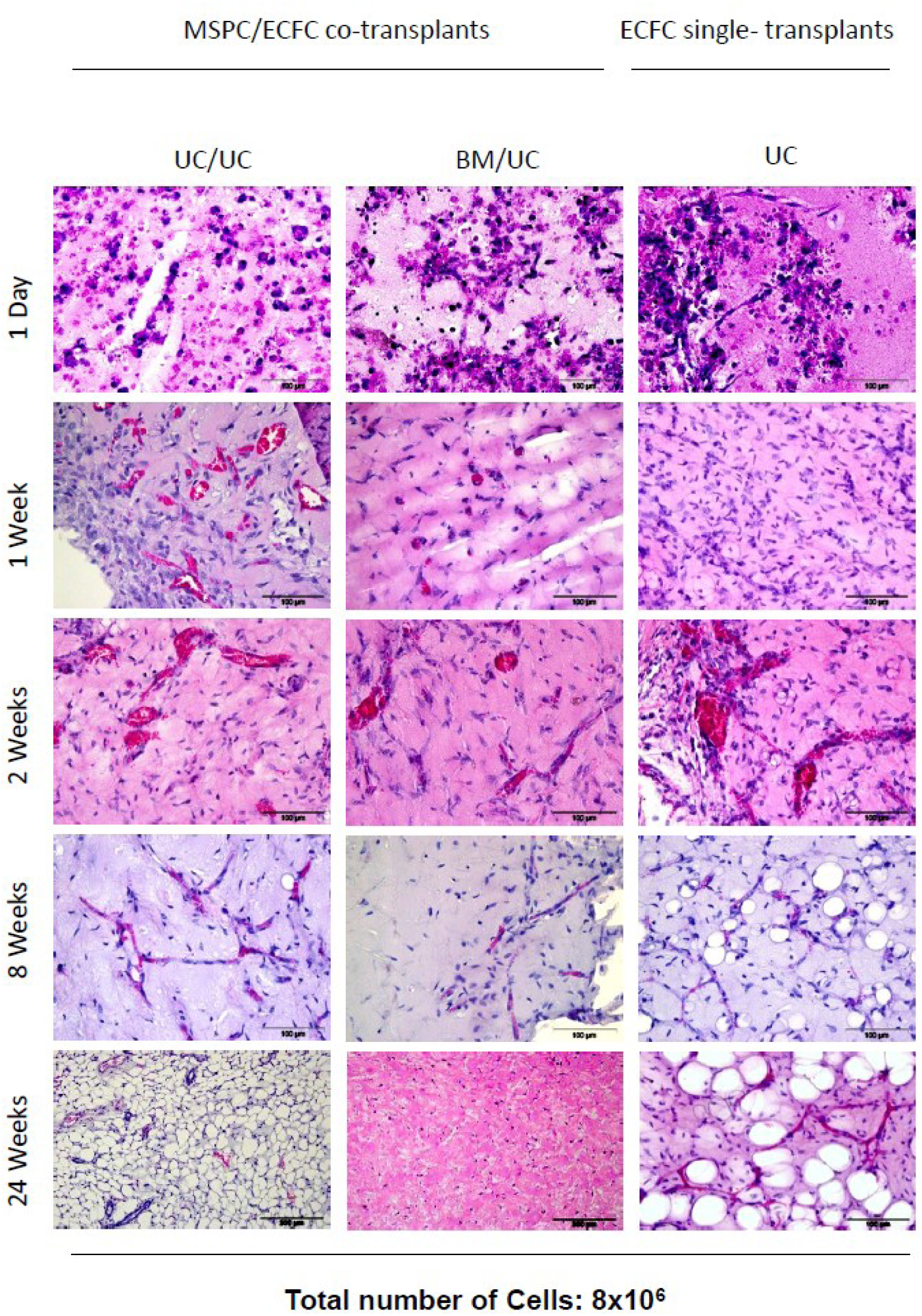

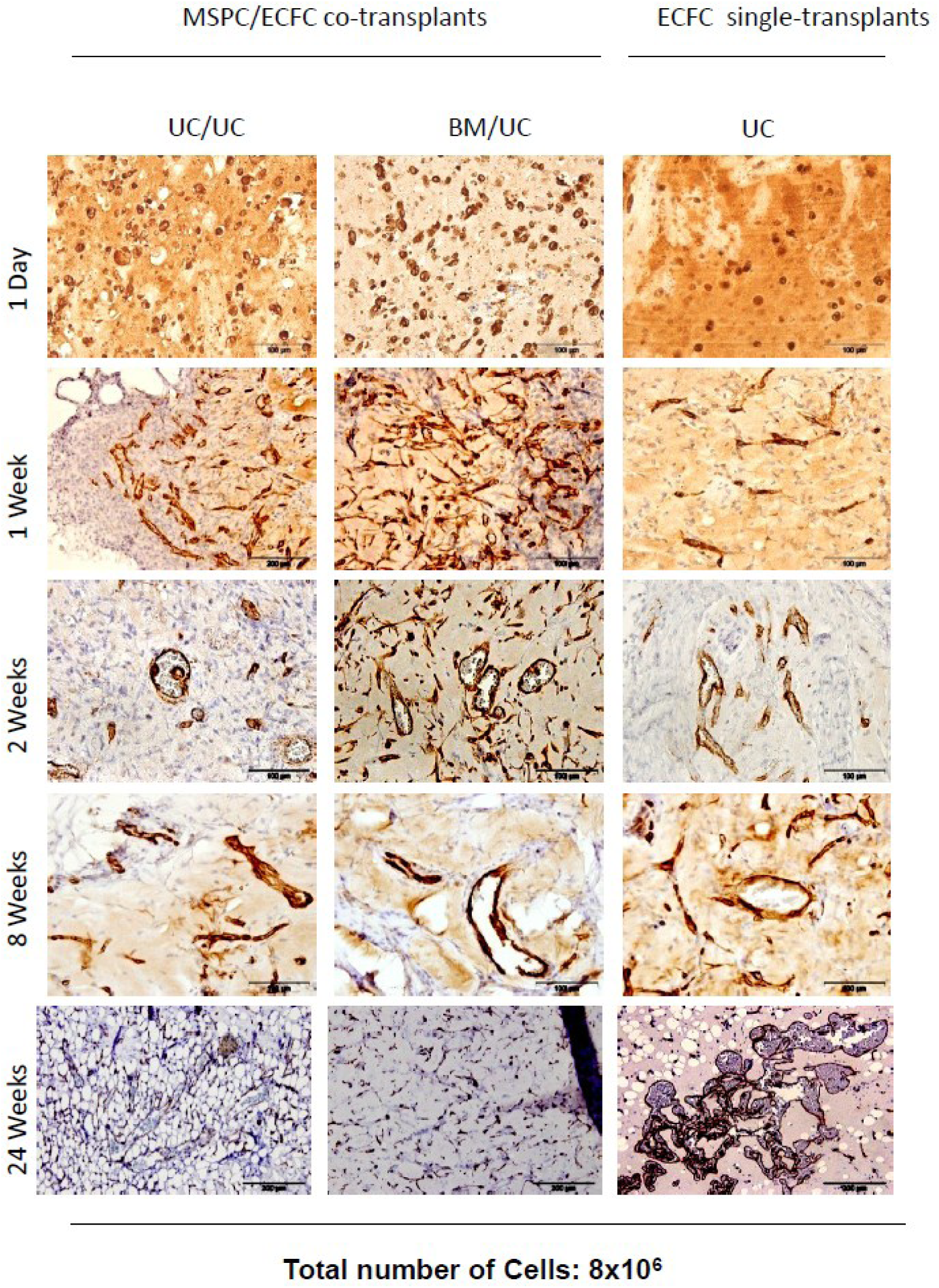

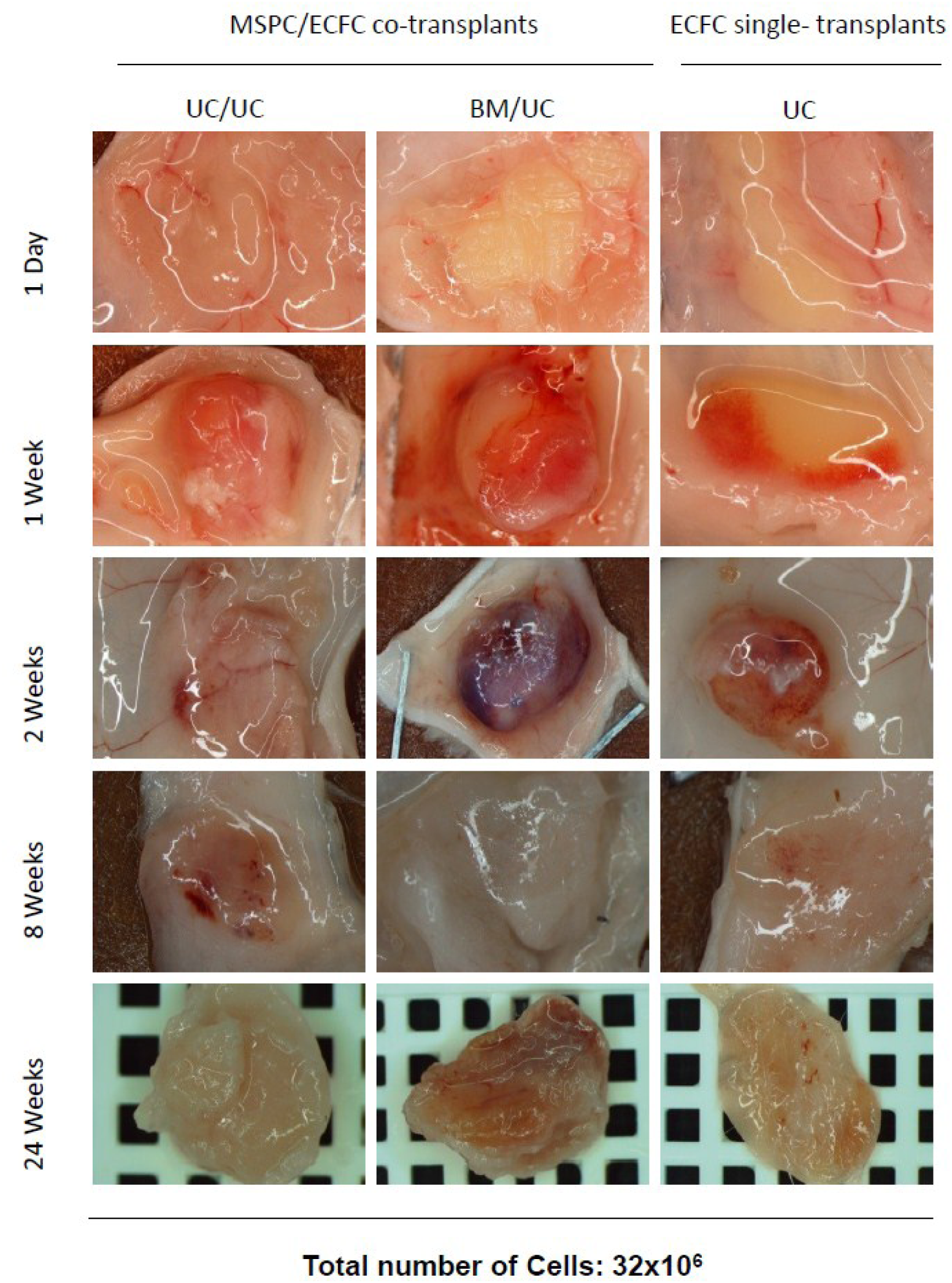

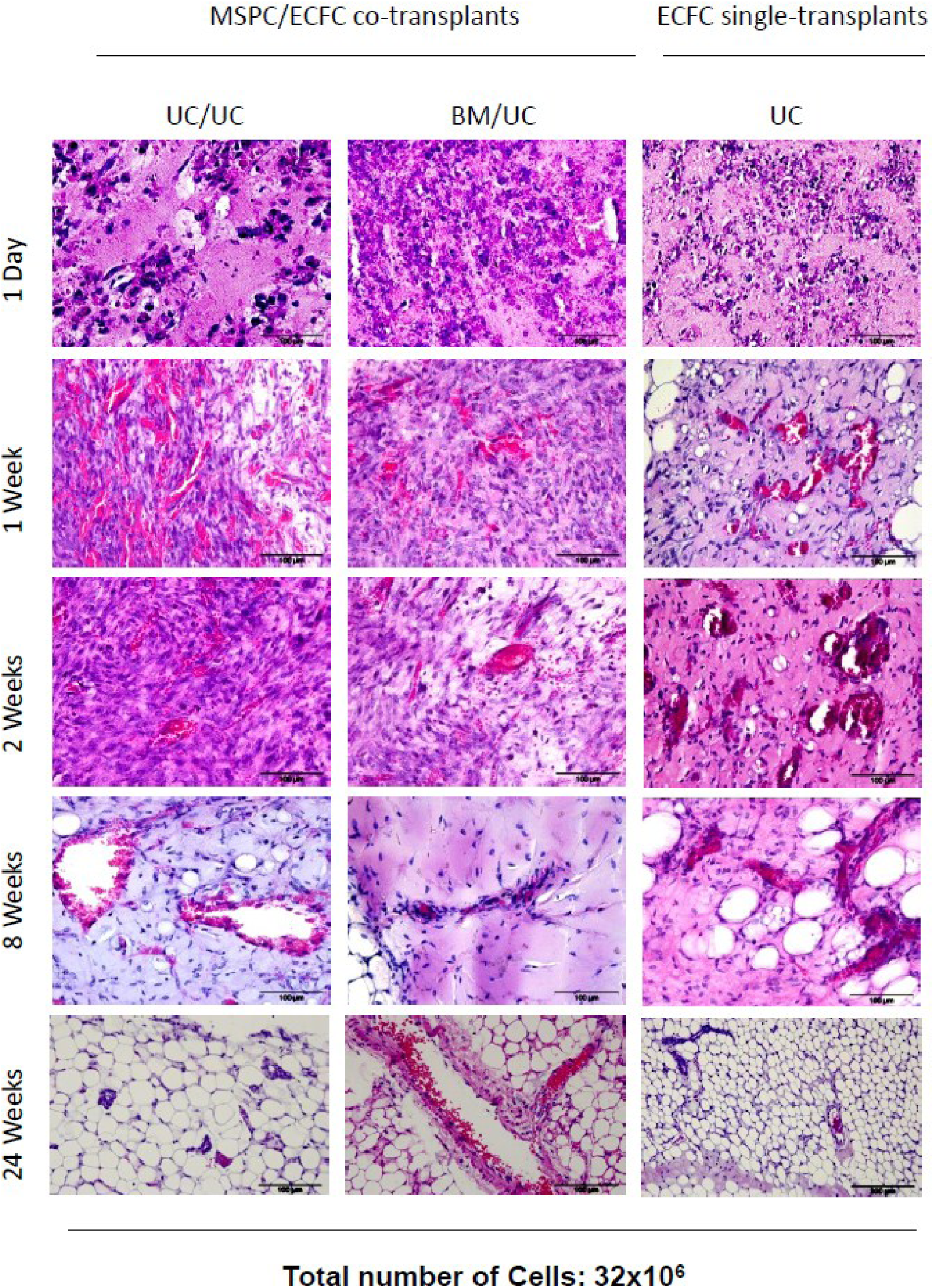

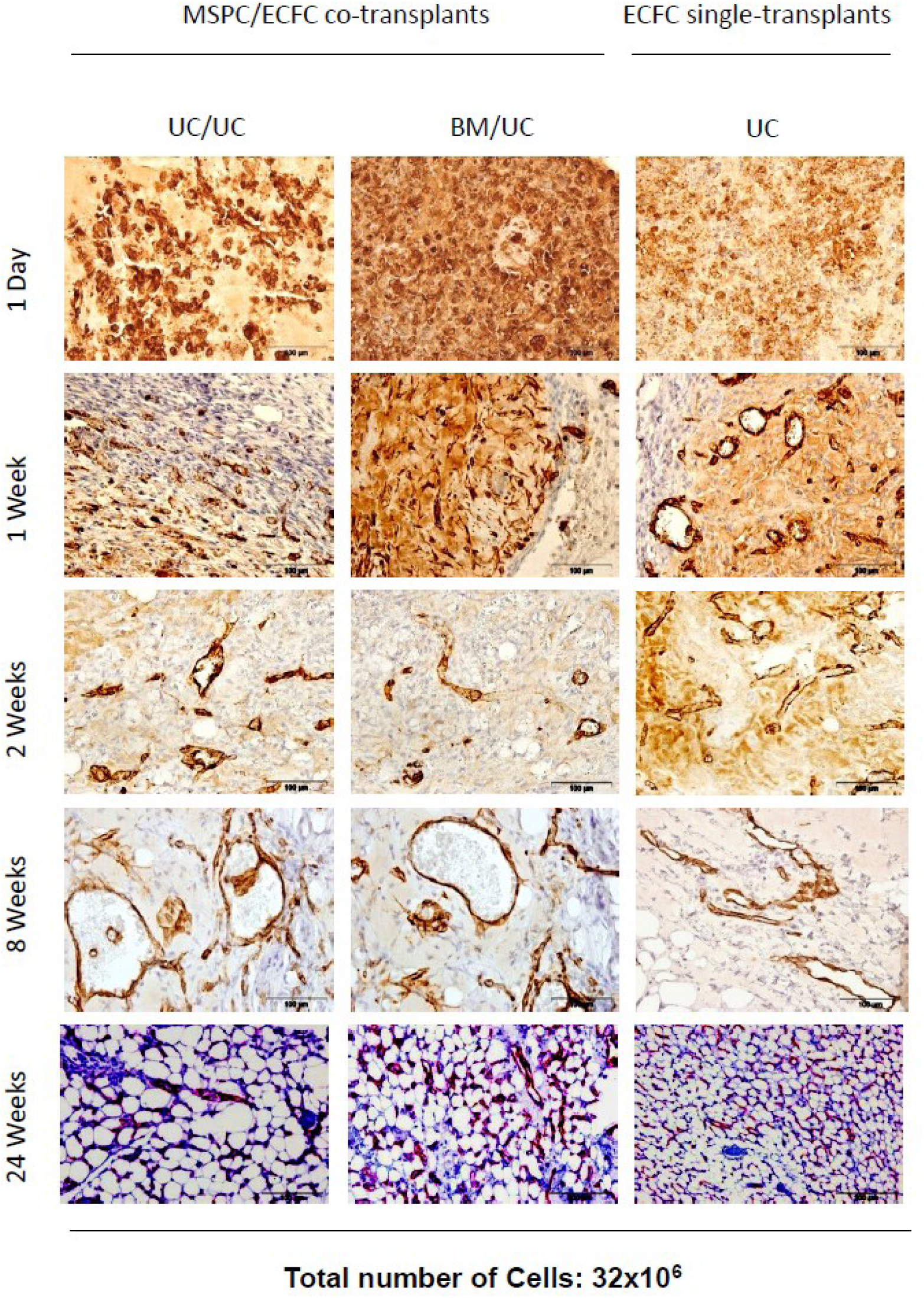

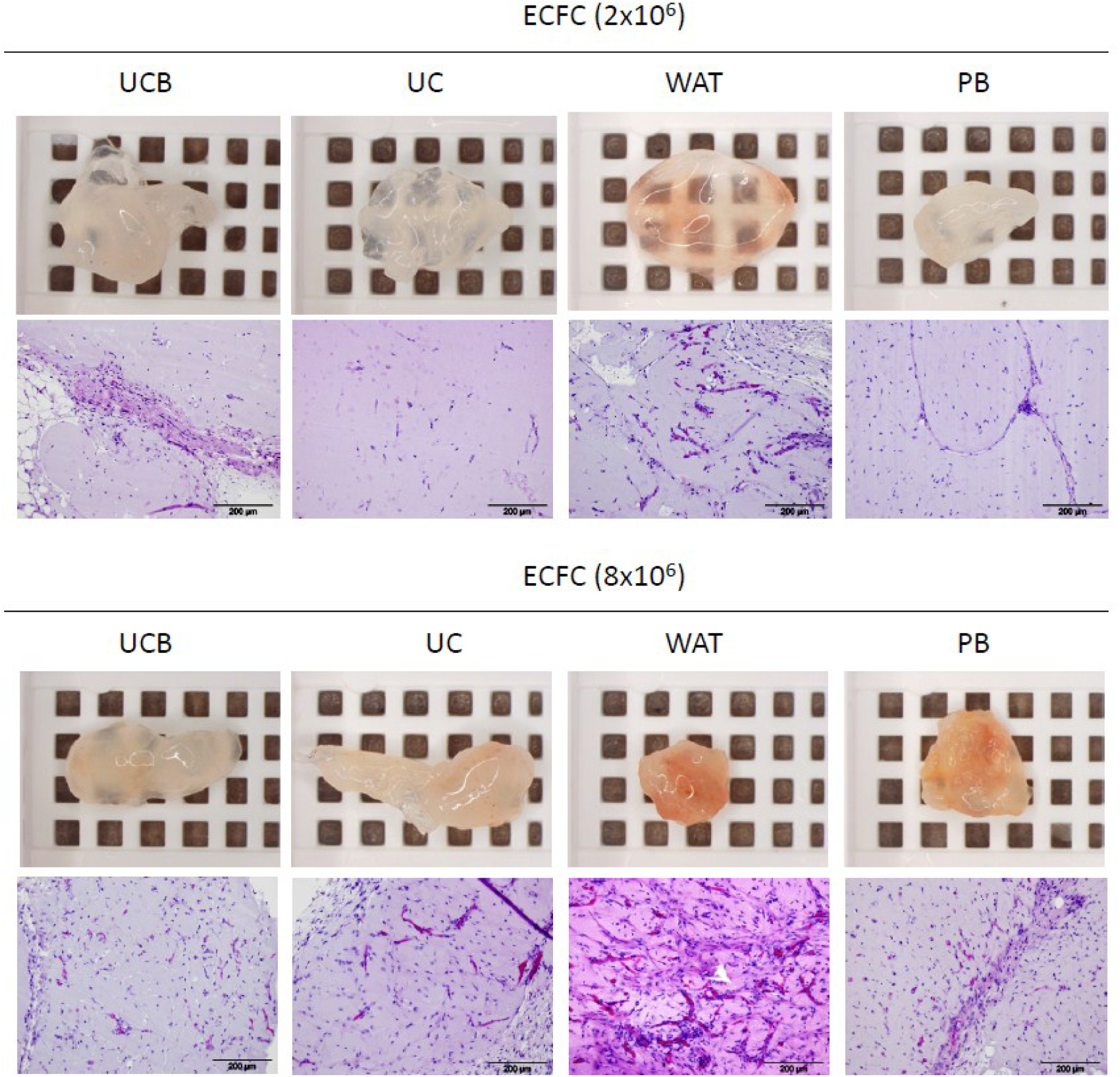

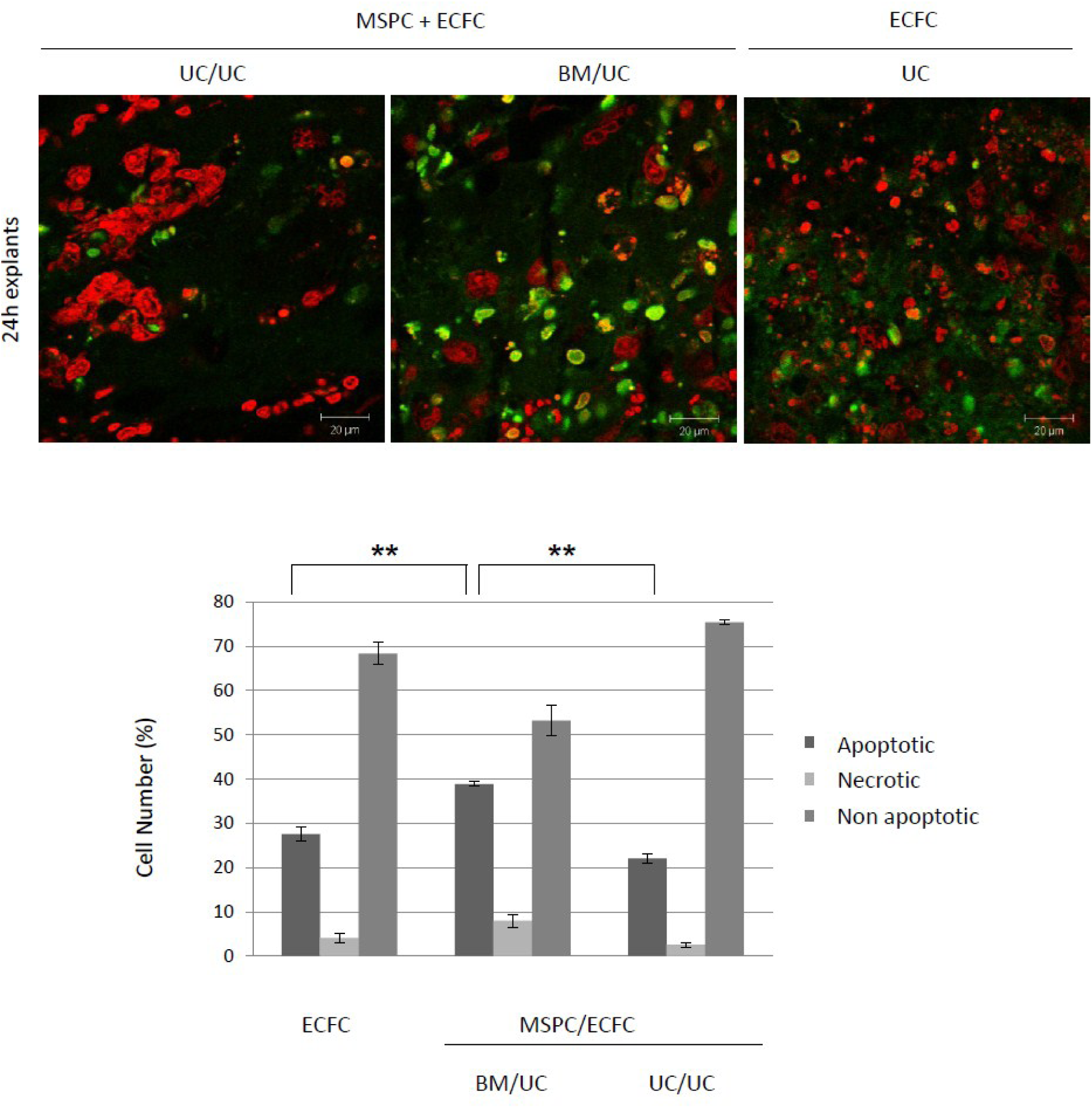

